# Effects of constitutive and acute Connexin 36 deficiency on brain-wide susceptibility to PTZ-induced neuronal hyperactivity

**DOI:** 10.1101/2020.07.27.223651

**Authors:** Alyssa A. Brunal, Kareem C. Clark, Manxiu Ma, Y. Albert Pan

## Abstract

Connexins are transmembrane proteins that form hemichannels allowing the exchange of molecules between the extracellular space and cell interior. Two hemichannels from adjacent cells dock and form a continuous gap junction pore, thereby permitting direct intercellular communication. Connexin 36 (Cx36), expressed primarily in neurons, is involved in the synchronous activity of neurons and may play a role in aberrant synchronous firing, as seen in seizures. To understand the reciprocal interactions between Cx36 and seizure-like neural activity, we examined three questions: a) does Cx36 deficiency affect seizure susceptibility, b) does seizure-like activity affect Cx36 expression patterns, and c) does acute blockade of Cx36 conductance increase seizure susceptibility. We utilize the zebrafish pentylenetetrazol (PTZ; a GABA(A) receptor antagonist) induced seizure model, taking advantage of the compact size and optical translucency of the larval zebrafish brain to assess how PTZ affects brain-wide neuronal activity and Cx36 protein expression. We exposed wild-type and genetic Cx36-deficient (*cx35*.*5- /-*) zebrafish larvae to PTZ and subsequently mapped neuronal activity across the whole brain, using phosphorylated extracellular-signal-regulated kinase (pERK) as a proxy for neuronal activity. We found that *cx35*.*5-/-* fish exhibited region-specific susceptibility and resistance to PTZ-induced hyperactivity compared to wild-type controls, suggesting that genetic Cx36 deficiency may affect seizure susceptibility in a region-specific manner. Regions that showed increased PTZ sensitivity include the dorsal telencephalon, which is implicated in human epilepsy, and the lateral hypothalamus, which has been underexplored. We also found that PTZ-induced neuronal hyperactivity resulted in a rapid reduction of Cx36 protein levels. 30 minutes and one-hour exposure to 20 mM PTZ significantly reduced the expression of Cx36. This Cx36 reduction persists after one-hour of recovery but recovered after 3-6 hours. This acute downregulation of Cx36 by PTZ is likely maladaptive, as acute pharmacological blockade of Cx36 by mefloquine results in increased susceptibility to PTZ-induced neuronal hyperactivity. Together, these results demonstrate a reciprocal relationship between Cx36 and seizure-associated neuronal hyperactivity: Cx36 deficiency contributes region-specific susceptibility to neuronal hyperactivity, while neuronal hyperactivity-induced downregulation of Cx36 may increase the risk of future epileptic events.

## INTRODUCTION

Connexins are transmembrane proteins that oligomerize to form a transmembrane pore called a hemichannel, which enables the exchange of molecules between the extracellular space and cell interior. Two hemichannels between adjacent cells can dock and form a continuous pore, known as a gap junction, allowing for direct intercellular coupling. Inter-neuronal gap junctions form electrical synapses, which are responsible for fast synaptic transmission and the synchronous firing of neurons within the brain (Rash et al., 2012). Connexin 36 (Cx36) is the main connexin expressed by neurons. It is involved in brain functions that rely on synchronous firing such as learning and memory (Allen, Fuchs, Jaschonek, Bannerman, & Monyer, 2011; Wang & Belousov, 2011), retina visual processing (Kovács-Öller et al., 2017), and sensorimotor reflex in the zebrafish (Miller et al., 2017). As the key structural component of electrical synapses, Cx36 may also act as a therapeutic target in diseases involving deficiencies in fast communication and aberrant synchronous firing, such as seizures. However, the reciprocal relationships between the Cx36 and seizures have remained unclear.

Previous work has examined the roles of Cx36 in the pathogenesis of seizures, but there has been no consensus on whether Cx36 increases or decreases seizure susceptibility (Gajda, Szupera, Blazsó, & Szente, 2005; Jacobson et al., 2010; Shin, 2013; Voss, Mutsaerts, & Sleigh, 2010a). Jacobson et al. (2010) found that Cx36 knockout mice exhibited an increase in seizure-like behaviors following the administration pentylenetetrazol (PTZ; a GABA(A)-receptor antagonist), indicating that normal expression of Cx36 may be protective against seizure-inducing conditions. However, this finding contradicts studies using the connexin blocking drug quinine, which found the drug either decreased the severity of seizures (Gajda et al., 2005), or showed no change (Voss, Mutsaerts, & Sleigh, 2010b). The discrepancy may potentially be due to the difference between chronic Cx36 deficiency (Cx36 knockout) versus acute Cx36 deficiency (quinine). However, quinine has broad antagonistic activity against many different connexins expressed in the nervous system, and the effects cannot be attributed solely to the inhibition of Cx36 (Cruikshank et al., 2004; Manjarrez-Marmolejo & Franco-Pérez, 2016). Additionally, the difference in seizure induction methods and seizure metrics also makes direct comparisons between studies problematic.

Previous findings are also mixed regarding how neuronal hyperactivity affects the expression of Cx36. In rodent seizure models and epilepsy patient post-mortem samples, some groups have found that Cx36 expression was increased (Collignon et al., 2006; Laura, Xóchitl, Anne, & Alberto, 2015; X. Wu, Wang, Hao, & Feng, 2017), while others found decreased Cx36 expression (Condorelli, Trovato-Salinaro, Mudo, Mirone, & Belluardo, 2003; Söhl et al., 2000) or no change (Motaghi, Sayyah, Babapour, & Mahdian, 2017). Furthermore, even though seizures result in brain-wide changes in neuronal connectivity (Morgan, Gore, & Abou-Khalil, 2010), seizure-induced changes in Cx36 expression had only been examined in the dorsal telencephalon (cortex and hippocampus) (Condorelli et al., 2003; Laura et al., 2015; Motaghi et al., 2017; X. L. Wu et al., 2018). Potential changes to Cx36 expression in other brain areas following neuronal hyperactivity remain unknown.

To further investigate the relationship between Cx36 and neuronal hyperactivity and address the technical limitations listed above, we employ zebrafish as an experimental system. The small size of zebrafish larvae facilitates imaging of the whole brain under a laser scanning confocal microscope, which provides a unique opportunity to examine whole-brain activity as well as dynamic Cx36 protein regulation in an intact vertebrate organism. Additionally, the PTZ-induced seizure model in zebrafish has been well-characterized physiologically and behaviorally (Afrikanova et al., 2013; S.C. Baraban, Taylor, Castro, & Baier, 2005; Burrows et al., 2020; Copmans, Siekierska, & de Witte, 2017) and is an effective model in identifying therapeutics to target epilepsy in humans (Scott C. Baraban, Dinday, & Hortopan, 2013).

Using zebrafish, we created a whole-brain activity map following hyperactivity using the MAP-mapping method (Randlett et al., 2015) to determine that there are both dose-varying and region-specific changes in neuronal hyperactivity following administration of PTZ. Additionally, we created a whole-brain expression map of Cx36 following the administration of PTZ. With this, we determined specific brain regions that showed decreases in Cx36 expression following hyperactivity. Finally, by acutely reducing the function of Cx36 using the Cx36 blocking drug, mefloquine, we determined that acute inhibition of Cx36 is detrimental, and leaves the animal more susceptible to PTZ-induced hyperactivity than their untreated counterparts.

## METHODS

### Zebrafish Husbandry

All zebrafish used in this study were pigmentless (*nacre-/-*) in a mixed background of AB and TL wild-type strains (Zebrafish International Resource Center). *cx35*.*5* (ZFIN gene symbol: *gjd2a*) heterozygotes were gifts from Dr. Adam Miller at the University of Oregon (Marsh, Michel, Adke, Heckman, & Miller, 2017). Zebrafish embryos and larvae were raised under 14 h light/10 h dark cycle at 28.5°C in water containing 0.1% Methylene Blue hydrate (Sigma-Aldrich). Sex is not a relevant variable for the larval stages being used (0-6 days post-fertilization, dpf), as laboratory zebrafish remain sexually undifferentiated until two weeks of age (Maack & Segner, 2003; Wilson et al., 2014). All husbandry procedures and experiments were performed according to protocols approved by the Institutional Animal Care and Use Committee at Virginia Tech.

### Immunohistochemistry

Zebrafish larvae were fixed overnight in 4% paraformaldehyde (PFA) on a rocker at 4°C. Samples were then processed and stained as previously described by Randlett et al. (2015). Primary antibodies that were used are as follows: p44/42 MAPK (tERK) (L34F12, Cell Signaling Technologies), Phospho-p44/42 MAPK (pERK) (D13.14.4E, Cell Signaling Technologies), and Anti-activated Caspase 3 (BD Pharmingen). For the Connexin antibody (36/GJA9, Invitrogen), fish were fixed in 2% TCA for 3 hours, and sample processing and staining were performed as previously described (Marsh et al., 2017).

### MAP-(Activity Map)

Wild-type and *cx35*.*5* mutant in 6 dpf zebrafish larvae were first acclimated for 15 minutes in a 6-well plate and then transferred into a well containing 0 mM (E3 embryo media only), 2 mM, 5 mM, 10 mM, or 20 mM PTZ in embryo media for 15 minutes. Larvae were then fixed in 4% PFA overnight and immunostained and imaged using a Nikon A1 confocal microscope. Subsequent MAP-mapping analysis was performed as previously described (Randlett et al., 2015). Brain regions highlighted in the text of this document were selected based on the following criteria: only brain regions were selected (individual neuron clusters were not mentioned), and only brain regions with well-defined functions were selected to be highlighted. All identified brain regions and neuron clusters can be found in the Supplementary tables.

### Cx36 Expression Map

6 dpf larvae were acclimated for 15 minutes in a 6-well plate with embryo media and then transferred into a well containing 20 mM PTZ for either 30 minutes or 1 hour. Larvae were then either fixed immediately or allowed to recover for 1 hour, 3 hours, 6 hours, or 24 hours in embryo media. Larvae were fixed in 2% trichloroacetic acid (TCA) for 3 hours and immunostained as previously described (Miller et al., 2017). Confocal images were then morphed to a tERK standard brain image stack using CMTK (Randlett et al., 2015). To subtract background signal, an average stack of *cx35*.*5-/-* fish morphed and stained in the same way was subtracted from all images and then were processed as previously described, except for replacing pERK with the morphed and background subtracted anti-Cx36 (Randlett et al., 2015).

### Cell Death Quantification

6 dpf mutant and wild-type larvae were first acclimated for 15 minutes in a 6-well plate and then transferred into a well containing either embryo medium or 20 mM PTZ for 1 hour. Larvae were then immediately fixed in 4% PFA overnight, and immunostained Images were morphed to a standard brain and analyzed as previously described (Randlett et al., 2015). ROIs for the Diencephalon, Mesencephalon, Telencephalon from ZBrain were then overlaid on each stack, and Caspase positive cells were counted in each ROI. Standard unpaired t-tests with Welch’s correction for multiple comparisons were run between each group in GraphPad Prism.

### Mefloquine Treatment

At 6 dpf, larvae were exposed to either 0.025% DMSO (vehicle group) or 25 μM mefloquine. After 3 hours of exposure, fish and their relative media (either DMSO or mefloquine) were transferred to a 6 well plate and allowed to acclimate for 15 minutes. Larvae were then transferred to embryo media with 0 mM, 2 mM, 5 mM, 10 mM, or 20 mM PTZ for 15 minutes. Larvae were then immediately fixed in 4% PFA overnight, immunostained, and imaged using a Nikon A1 confocal microscope. Subsequent analysis was performed as previously described (Randlett et al., 2015).

### Image Processing and Statistical Analysis

Images were processed and quantified using Fiji (Schindelin et al., 2012). MATLAB 2019 (MathWorks) was used for MAP-mapping analysis (Randlett et al., 2015). For Caspase-3 quantification, statistical analyses were performed in GraphPad Prism (Version 8). Student’s t-test with Welch’s correction for multiple comparisions was preformed. Results were considered significant if *p*<0.05.

## RESULTS

### PTZ induces brain-wide neuronal hyperactivation in a dose-dependent manner

PTZ inhibits GABA(A) receptor-mediated inhibitory neurotransmission, which leads to global neuronal hyperactivation and seizure-like neurological and behavioral phenotypes in both rodents and zebrafish (S.C. Baraban et al., 2005). To determine whether different brain regions have distinct sensitivities to PTZ-induced neuronal hyperactivation, we first compared whole-brain activity maps in wild-type fish exposed to varying concentrations of PTZ. To do this, we utilized the MAP-mapping assay to create whole-brain activity maps (Randlett et al. 2015). MAP-mapping utilizes the ratio of total extracellular signal-regulated kinase (tERK), which is present in all neurons, and phosphorylated ERK (pERK), the phosphorylated form of ERK that is induced (within 10 minutes) following neuronal activity. The ratiometric pERK/tERK signal can then be quantified and statistically tested in an annotated 3D brain atlas (Z-Brain) (Randeltt et al. 2015).

Using MAP-mapping, we found region-specific changes in neuronal activity in response to varying concentrations of PTZ. We treated wild-type animals by bath-exposing them to embryo media with 2, 5, 10, and 20 mM PTZ for 15 minutes. Animals exposed to media only were used as the baseline for comparison. Neuronal activity was measured by the pERK/tERK ratio as described previously (Randlett et al., 2015). After exposure to 2 mM PTZ, we saw moderate increases in neuronal activity in more restricted brain areas in regions responsible for homeostatic regulation (hypothalamus and preoptic area) and executive functioning (subpallium, pallium) as well as the cerebellum (Figure 1A). After exposure to 5, 10, and 20 mM PTZ, we observed broader increases in brain-wide neuronal activity (Figure 1B-D). These regions include those that were activated by 2 mM PTZ (hypothalamus, preoptic area, subpallium, and in many regions involved in movement control such as the pretectum, cerebellum, and oculomotor nuclei. Additionally, we observed some brain areas that became less active after exposure to PTZ: the telencephalon was less active at 10 and 20 mM PTZ than at lower concentrations (Figure 1D) and the olfactory bulb was less active across all PTZ concentrations (Figure 1A-D). The complete list of all identified changes is provided in Supplementary Table 1.

**Figure 1.**
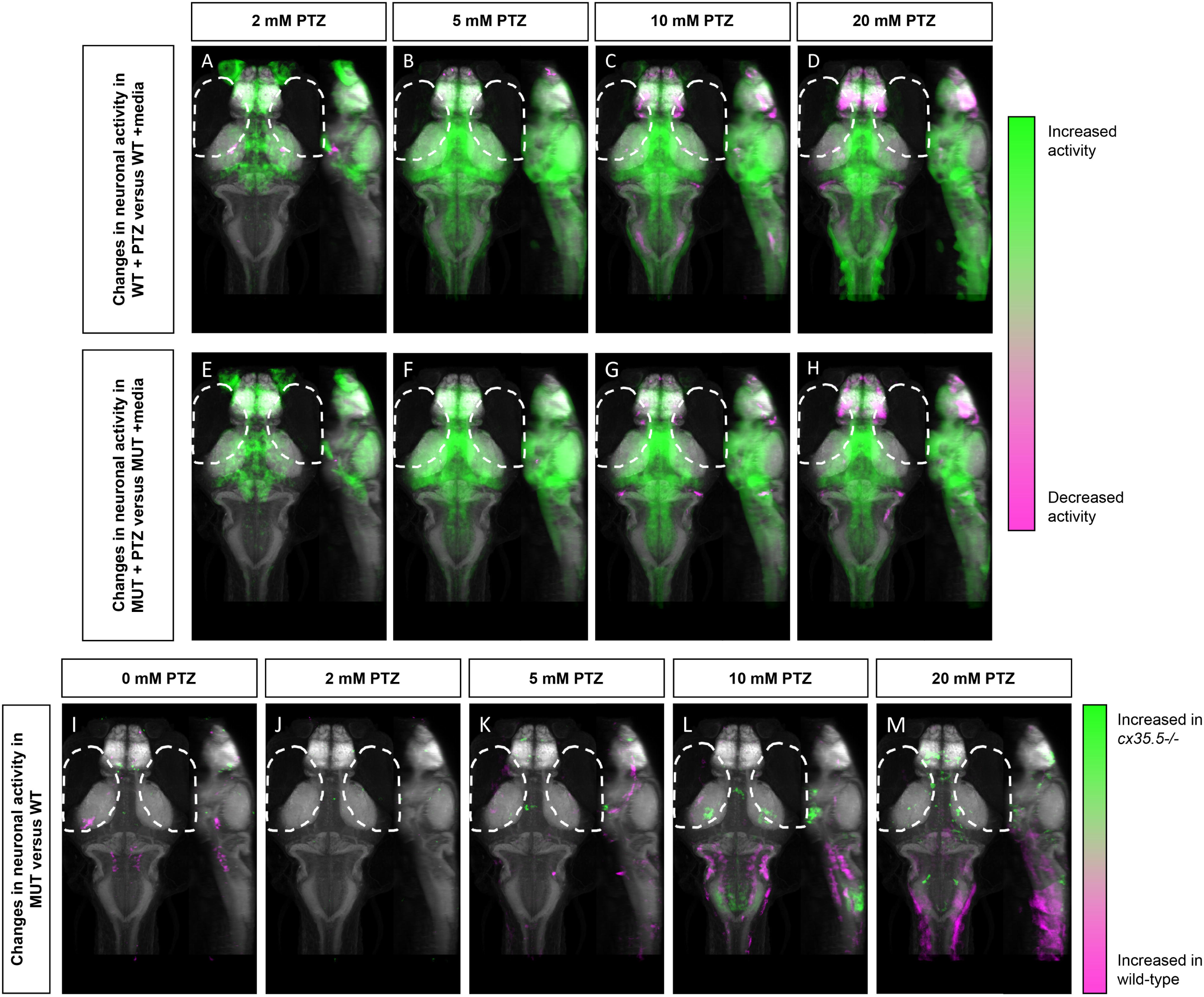
Whole-brain activity map showing significant regional differences in neuronal activity following various PTZ concentration exposure in wild-type and *cx35*.*5-/-* zebrafish larvae. Dorsal and lateral view of zebrafish larvae brain. Colors indicate ROIs with higher pERK/tERK ratio in wild-type PTZ treated (green) or in Embryo Media (magenta) in A) 2 mM PTZ treated (n=10) B) 5 mM PTZ treated (n=8) C) 10 mM PTZ treated (n=10) and D) 20 mM PTZ treated (n=10) vs Embryo Media (n=10). Colors indicate ROIs with higher pERK/tERK ratio in *cx35*.5 *-/-* larvae PTZ treated (green) or in Embryo Media (magenta) in E) 2 mM PTZ treated (n=9) F) 5mM PTZ treated (n=10) G) 10mM PTZ treated (n=11) and H) 20mM PTZ treated (n=10) vs Embryo Media (n=9). Colors indicate ROIs with higher pERK/tERK ratio in *cx35*.*5-/-* (green) or wild type (magenta) in I) E3 treated (WT n=10, MUT n=9) J) 2 mM PTZ treated (WT n=10, MUT n=9) K) 5 mM PTZ treated (WT n=8, MUT=10) L) 10 mM PTZ treated (WT n=10, MUT n=11) and M) 20 mM PTZ treated (WT n=10, MUT n=10) *cx35*.*5-/-* vs WT Images show pixels with significantly increased pERK/tERK ratio for treated fish (green) and untreated fish (magenta). For all regions p < 0.005

Overall, we were able to generate a PTZ dose-varying whole-brain activity map in 6 dpf zebrafish. We saw increased neuronal activity in areas previously identified to be involved in PTZ induced hyperactivity such as the pallium and optic tectum (Liu & Baraban, 2019). We also identified additional regions that were previously unidentified such as the hypothalamus.

### Genetic Cx36 deficiency results in changes in PTZ-induced brain-wide neuronal hyperactivity

To understand what effect loss of Cx36 has on hyperactivity we examined whole-brain activity changes at different concentrations of PTZ in the *cx35*.*5-/-* larvae. As the expression of the two zebrafish paralogs of Cx36, Cx35.5 and Cx34.1, are mutually dependent, loss of *cx35*.*5* results in near-complete loss of both Cx36 paralogs (Miller et al., 2017 and also Figure 3B). We again employed the MAP-mapping technique to determine which brain regions show a significant difference between PTZ-treated mutants and untreated mutants. Similar to their wild-type siblings, at 2 mM PTZ, significant increases in neuronal activity in the preoptic area, subpallium, and the hypothalamus were observed (Figure 1E). Additionally, we saw increases in the retinal arborization fields associated with visual processing (Figure 1E). At 5, 10, and 20 mM PTZ, we found a very similar map to that of their wild-type siblings, with increases and decreases in many of the same major brain regions listed previously (Figure 1F-H). For a complete list of significantly change brain regions, see Supplementary Table 1.

**Figure 2.**
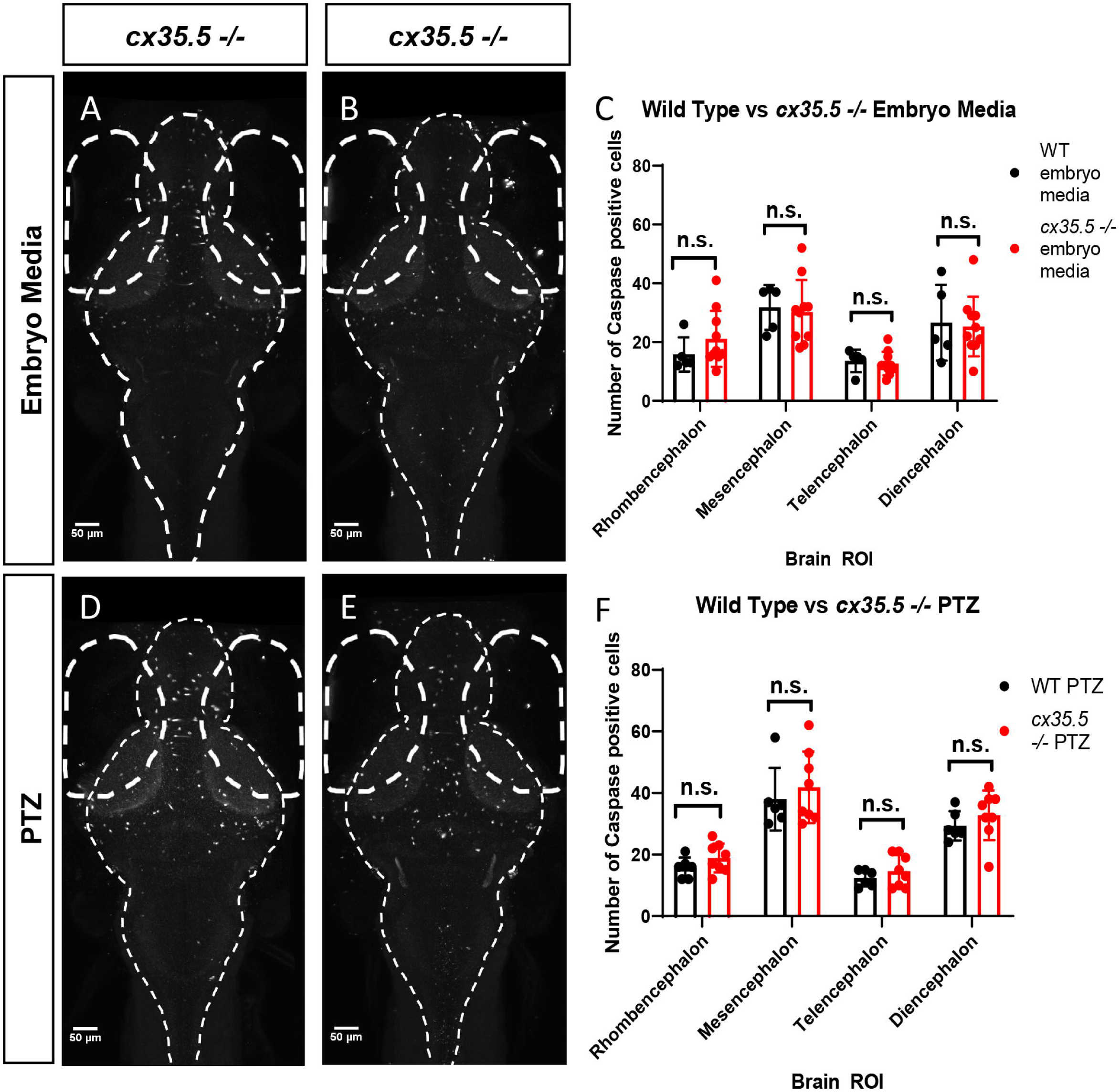
Caspase positive cells by major brain division, comparing *cx35*.*5-/-* vs wild type with and without PTZ. Sum stack projections of all fish (Caspase-3 staining) A-D. A graph depicting the number of Caspase-3 positive cells in the Rhombencephalon, Mesencephalon, Telencephalon, and Diencephalon in wild-type (black) vs *cx35*.*5-/-* (red) fish with treatment with E) Embryo medium (Vehicle) or F) PTZ. Data were analyzed using a student’s t-test with Welch’s correction. Embryo medium (vehicle) treatment, wild type n=5, *cx35*.*5-/-* n= 10. PTZ treatment, wild type n= 6 *cx35*.*5-/-* n=8.

**Figure 3.**
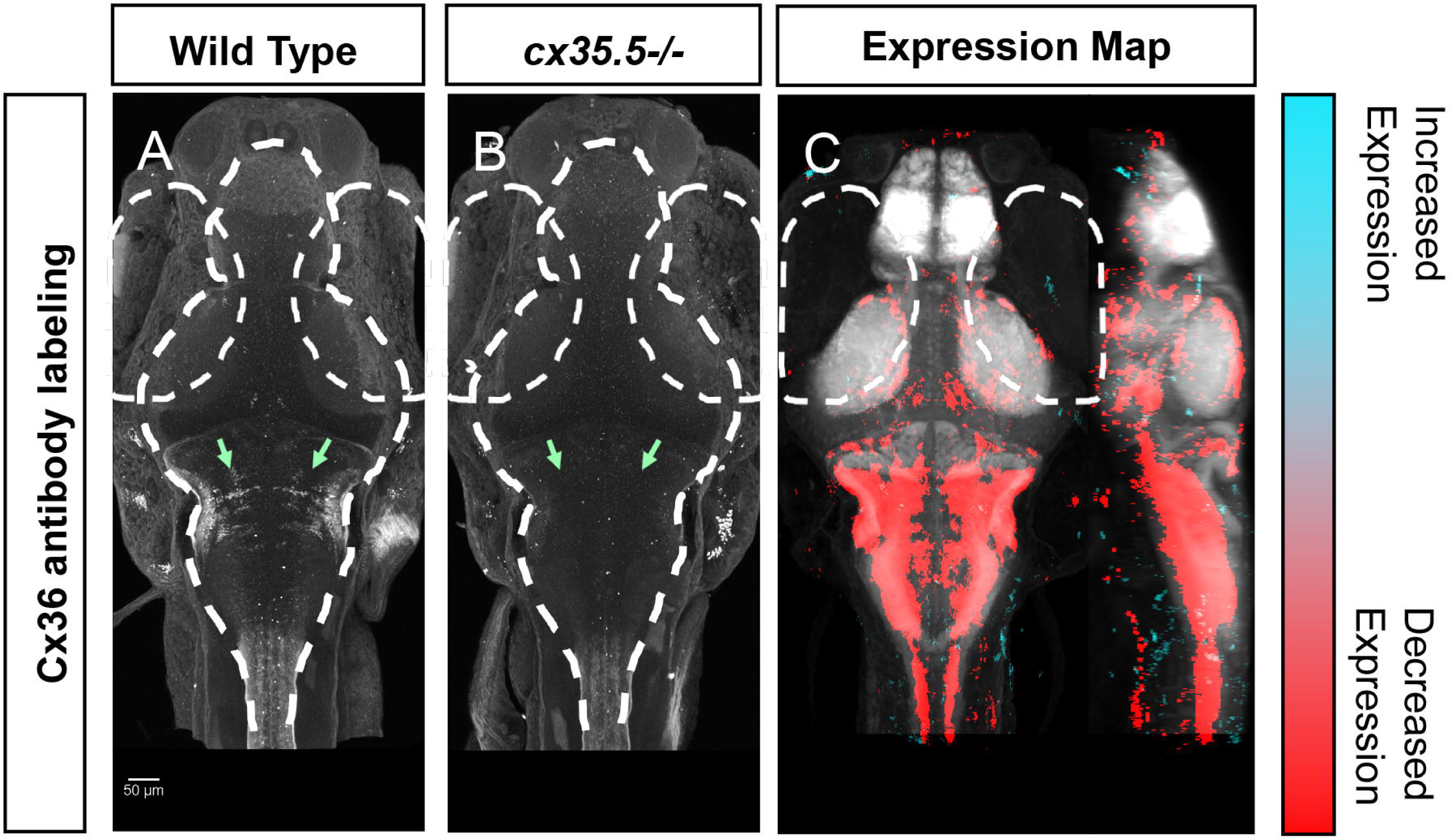
Whole-brain expression map of *cx35*.*5-/-* vs wild-type zebrafish larvae immunostaining of anti-human Cx36. Whole-brain expression of Cx36 using an anti-human Cx36 antibody vs tERK. Cyan indicates increases in fluorescence over tERK in *cx35*.*5-/-* fish, red indicates increases in fluorescence over tERK in wild-type fish A) Cx36 immunostaining of *cx35*.*5 -/-* fish B) Cx36 immunostaining of wild-type fish C) Dorsal and lateral view of zebrafish larvae brain. Whole-brain expression map showing increased expression in *cx35*.*5-/-* (cyan) and increased expression in wild type (red). Wild type n=10, *cx35*.*5-/-* n=7 p<0.005)

### Changes in *cx35*.*5-/-* whole-brain activity maps compared to wild-type

To understand differences in neuronal hyperactivity between *cx35*.*5 -/-* and wild-type animals, we compared the activity map of *cx35*.*5-/-* and wild-type siblings at baseline (media only) and after exposure to different concentrations of PTZ (Figure 1I-M). We observed no increases in neuronal activity at baseline, however, we did observe decreases in activity in *cx35*.*5-/-* relative to wild-type in the rhombencephalon reticulospinal neurons and medial vestibular neurons (Figure 1I). At 2 mM PTZ, there were no significant changes in brain-wide neuronal activity between *cx35*.*5-/-* and wild-type siblings (Figure 1J). At 5 mM PTZ, there were small increases in activity in the hypothalamus and the subpallium (Figure 1K). At 10 mM PTZ, we observed increases in the hypothalamus and various regions within the rhombencephalon (Figure 1L). We also found regions that show less of an increase in activity in *cx35*.*5-/-* compared to wild-type within the rhombencephalon specifically in regions that rely on the synchronous firing capabilities of Cx36 (Mauthner cells, inferior olive) (Bazzigaluppi et al., 2017; Flores et al., 2012; Yao et al., 2014). At the highest concentration (20 mM), we saw increased activity in the *cx35*.*5-/-* compared to wild-type in areas previously identified as associated with seizures such as the pallium (Liu & Baraban, 2019) as well as the hypothalamus. These regions are similar to our findings in the wild-type animals after PTZ exposure, indicating an increase in severity of hyperactivity in these regions following treatment with PTZ in *cx35*.*5-/-* animals. We also observed regions that show fewer increases in activity in the rhombencephalon, relative to wild-type, similar to 10 mM PTZ, but they are less severe (Figure 1L, M). For a complete list of regional differences, please see Supplementary Table 1.

### Genetic Cx36 deficiency does not affect cell death at baseline or after PTZ

We determined that PTZ alone and PTZ in combination with *cx35*.*5* deficiency resulted in regional and dose-varying changes in whole-brain neuronal activity. One possible explanation is that *cx35*.*5* mutation may result in altered neuronal cell death, either at baseline or after PTZ, which would then alter the overall balance of brain-wide connectivity. To test this, we stained for activated caspase-3 (a marker of apoptotic cells) and quantified the number of positive cells in each of the major brain divisions (rhombencephalon, mesencephalon, telencephalon, and diencephalon). We found that there were no differences at baseline (media only) in the number of caspase-3 positive cells between *cx35*.*5-/-* and wild-type siblings in any of the major brain divisions (Figure 2A, B, D, E). Additionally, no difference in the number of caspase-3 positive cells when comparing both *cx35*.*5-/-* and wild-type siblings after 20 mM PTZ was found (Figure 2C, F). From these data, we, therefore, conclude that changes in neuronal response in *cx35*.*5* animals are not likely caused by altered cell death induction.

### Creation of the whole-brain Cx36 expression map

To understand how neuronal hyperactivity affects Cx36, we created a whole-brain *expression* map to efficiently, and in a non-biased manner, measure changes in protein expression using a modified MAP-mapping processing procedure. We utilized a previously-validated human anti-Cx36 antibody and stained wild-type (Figure 3A) and *cx35*.*5-/-* (Figure 3B) siblings. Consistent with previous studies, near-complete loss of anti-Cx36 staining in *cx35*.*5-/-* animals was detected. To quantify Cx36 expression across the whole brain, we performed image normalization (with CMTK) and subtracted the average stack of all *cx35*.*5-/-* fish from each animal. We then followed the same MAP-mapping processing pipeline to quantify the Cx36/tERK ratio, with tERK staining used to normalize staining intensity across animals. The resulting Cx36 expression map reveals decreases in Cx36 staining intensity in *cx35*.*5-/-* fish compared to wild-type siblings in regions such as the optic tectum, rhombomeres, mauthner cells, etc. (Figure 3C). See Supplementary Table 2 for a complete list of regional changes. We then applied this same method to examine Cx36 expression after PTZ.

### Reduced Cx36 expression following PTZ exposure

Next, to determine if exposure to PTZ changes Cx36 expression, we compared the Cx36 expression map between treated animals and untreated animals exposed to 20 mM PTZ for 30 minutes or 1 hour. After 30 minutes of PTZ exposure, we found a global decrease in Cx36 fluorescence (Figure 4A). A similar but more pronounced effect was observed after 1 hour (Figure 4B). We saw decreases in Cx36 expression in the optic tectum, the retinal arborization fields, and in the rhombencephalon in rhombomere 7, an area that is important for motor behavior (Figure 4A, B). After 1 hour of PTZ, there was also a decrease in expression within the cerebellum (Figure 4B), an area that relies heavily on Cx36 for synchronous firing. For a complete list of ROIs with changes, see Supplementary Table 2. Together, these data reveal that Cx36 expression is reduced following exposure to PTZ after 30 minutes, and this is exacerbated after 1 hour of exposure.

**Figure 4.**
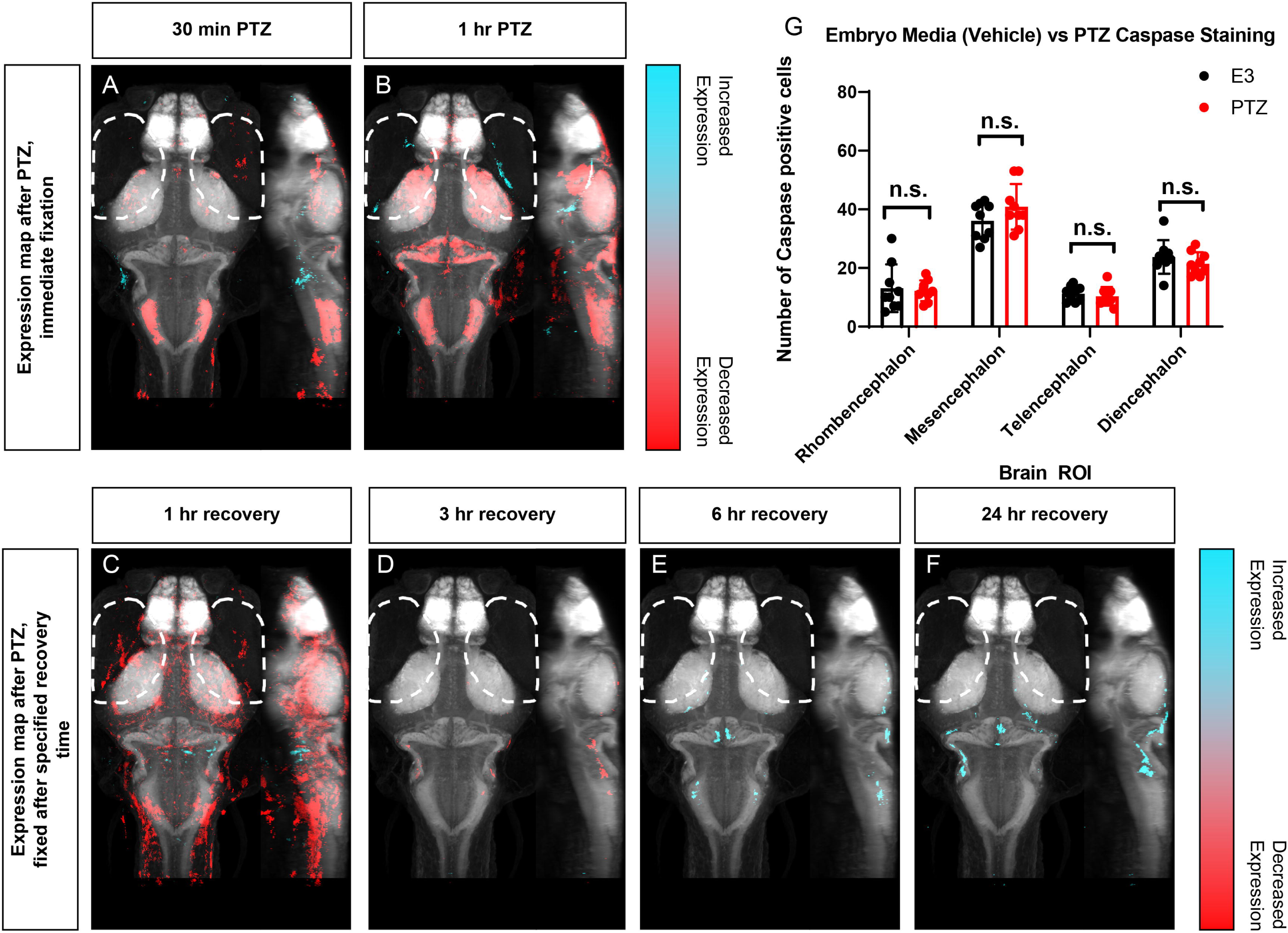
Wild type whole-brain immunostaining Cx36 expression map in E3 vs PTZ treated zebrafish larvae. Dorsal and lateral view of zebrafish larvae brain. Whole-brain expression of Cx36 using an anti-human Cx36 antibody vs tERK. Cyan indicates increases in fluorescence over tERK in PTZ treated fish, red indicates increases in fluorescence over tERK in E3 treated fish A) After 30 min of 20 mM PTZ exposure (n=10) B) After 1 hr of 20 mM PTZ exposure (n=10) C) 1 hour recovery after PTZ is removed, n=12 D) 3 hours of recovery after PTZ is removed, n=12 E) 6 hours of recovery after PTZ is removed, n=12 F) 24 hours of recovery after PTZ is removed, n=10. For all regions p < 0.005 G) A graph depicting the number of Caspase-3 positive cells in the Rhombencephalon, Mesencephalon, Telencephalon, and Diencephalon in wild-type fish with treatment with embryo medium (Vehicle) (Black) or PTZ (Red). Data were analyzed using a student’s t-test with Welch’s correction, n=9.

### Recovery of Cx36 expression following cessation of PTZ exposure

To test whether Cx36 expression recovers after the removal of PTZ, we created Cx36 expression maps for animals exposed to 20 mM PTZ for one hour and then allowed them to recover in embryo media for 1, 3, 6, or 24 hours after PTZ removal. Compared to animals not exposed to PTZ, Cx36 expression was still significantly decreased in the pallium, habenula, subpallium, and the pretectum after 1 hour of recovery, but there were some increases in expression in restricted areas in the rhombencephalon (Figure 4C). The decrease in Cx36 expression was almost entirely recovered after 3 hours (Figure 4D). Interestingly, expression is then slightly increased by 6 hours of recovery in the optic tectum, neuropil, and the cerebellum (Figure 4E). This is maintained 24 hours later (Figure 4F). For a complete list of regions that show changes in expression, see Supplementary Table 2. These alterations in expression were not due to cell death resulting from long-term PTZ exposure as no significant differences in the number of caspase-3 positive cells in between untreated (media only) versus those treated with 20 mM PTZ for one hour (Fig. 4G) we detected.

### Acute blockade of Cx36 increases neuronal hyperactivity following PTZ exposure

Given that PTZ-induced neuronal hyperactivity resulted in decreased Cx36 expression, we next tested whether the acute reduction of Cx36 contributes to further susceptibility to neuronal hyperactivation, i.e., whether PTZ-induced Cx36 reduction is maladaptive. To acutely inhibit Cx36 function, we utilized a Cx36-specific blocking drug, mefloquine, and examined changes in neuronal activity. The effects of mefloquine were assessed by comparing the activity maps of wild-type fish treated with DMSO (vehicle) or 25 μM mefloquine for 3 hours before the experiment, with or without varying concentrations of PTZ. Similar to our wild-type activity mapping (Figure 1A-D), we observed broad increases in neuronal activity in DMSO treated animals following exposure to PTZ in a dose-dependent manner (Figure 5A-D), but these increases were greater than our wild-type treated control (Figure 1A-D). At 2 mM PTZ, we saw increases in activity in the optic tectum, neuropil, cerebellum, pallium, and hypothalamus. There were also decreases in activity in the olfactory bulb (Figure 5A). At 5 mM PTZ, we found increases in activity in similar regions as well as the retinal arborization fields and decreases in the olfactory bulb (Figure 5B). At 10 mM we observed increases in similar regions, with greater increases seen in the hypothalamus, decreases in the olfactory system and, small decreases in the hypothalamus and pallium (Figure 5C). Finally, at 20 mM PTZ increases in neuronal activity in similar regions as the previous doses were observed, with the greatest increases seen in the hypothalamus. Decreases in the olfactory system, hypothalamus, and pallium (Figure 5D) were also observed. In fish treated with mefloquine, we found very similar overall patters as the DMSO treated fish (Figure 5A-D), but at each dose, we saw increases in the hypothalamus, preoptic area and subpallium, and fewer decreases within the forebrain (Figure 5E-H).

**Figure 5.**
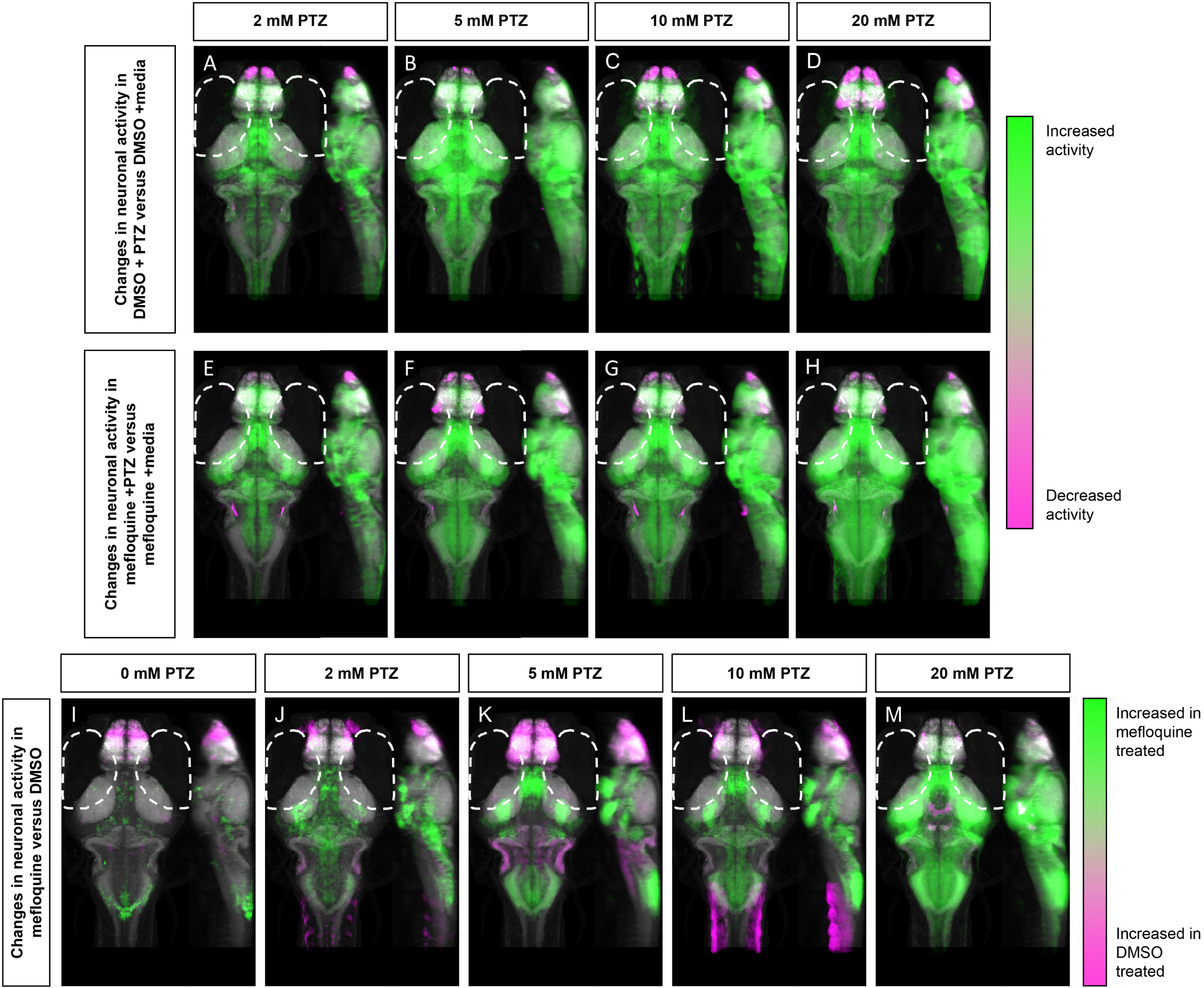
Whole-brain activity map showing significant regional differences following Connexin 36 blocking drug mefloquine and PTZ exposure in wild-type zebrafish larvae. Dorsal and lateral view of zebrafish larvae brain. Images show pixels with significantly increased pERK/tERK ratio compared to DMSO and embryo medium (n=9) for DMSO treated fish after A) 2 mM PTZ (n=10) (B) 5 mM PTZ (n=10) C) 10 mM PTZ (n=10) and D) 20 mM PTZ (n=10). E-H) Images show pixels with significantly increased pERK/tERK ratio compared to mefloquine and embryo medium treated fish (n=9) after E) 2 mM n=10) PTZ F) 5 mM PTZ (n=8) G) 10 mM PTZ (n=10) and H) 20 mM PTZ (n=10). I-M) Images show pixels with significantly increased pERK/tERK ratio for mefloquine (25 μM) treated fish (green) and for DMSO (Vehicle) treated fish (magenta) after exposure to I) embryo medium (n=9) J) 2mM PTZ (n=10) K) 5 mM (DMSO n=10, mefloquine n=8) PTZ L) 10 mM PTZ (n=10) M) 20 mM PTZ (n=10). For all regions p < 0.005

Next, we compared mefloquine versus DMSO treated siblings at different concentrations of PTZ. In the absence of PTZ, the mefloquine treated fish showed increases and decreases in neuronal activity in different brain regions, compared to DMSO treated siblings (Fig. 5I). Specifically, we saw moderate increases in the hypothalamus, cerebellum, and tegmentum. There were decreases in activity in the olfactory bulb and we observed less of an increase in activity compared to control in the telencephalon, specifically in the subpallium (Figure 5I). At 2 mM PTZ, mefloquine treated fish showed increases in the major regions associated with PTZ exposure (Figure 1A), compared to DMSO treated fish. Increases in the hypothalamus, retinal arborization fields, pre-tectum, and subpallium were found. There were decreases in the olfactory bulb and less of an increase in other regions of the telencephalon (Figure 5J) compared to control. At 5 mM PTZ, we found similar regions of increased activity in mefloquine treated fish, but we also saw regions that showed less of an increase in activity compared to control within both the telencephalon and the rhombencephalon (Figure 5K), specifically in regions that had high Cx36 expression (Figure 3C). At 10 and 20 mM PTZ, we observed similar increases in activity in mefloquine treated fish, each increasing with PTZ dose, and less of an increase in activity compared to control in the telencephalon, that was less severe than 10 mM, in these two groups (Figure 5L-M). At 20 mM we observed less of an increase in activity in the hypothalamus and oculomotor nuclei compared to wild-type, which was not observed in other doses (Figure 5M). The activity increases we found in the drug-treated animals are more wide-spread than in the *cx35*.*5 -/-* (Figure 1I-M), but similar regions were affected. These results indicate that acute reduction of Cx36 functionality results in increased susceptibility to PTZ-induced neuronal hyperactivity. For a complete list of regions changed, see Supplementary Table 3.

Finally, to test whether or not the increases in neuronal activity that we observed in mefloquine treated fish compared to *cx35*.*5-/-* fish were due to mefloquine’s off-target effects, we examined effects of mefloquine on *cx35*.*5-/-* fish, with and without PTZ. We compared the differences in neuronal activity in mefloquine treated and DMSO treated *cx35*.*5-/-* fish, with either no PTZ (embryo media only) or a moderate PTZ dose (5 mM PTZ) (Supplementary Figure 1). In both the embryo media and 5 mM PTZ conditions, we observed increases in neuronal activity following the administration of mefloquine in a small region of the rhombencephalon (area postrema, neuropil, rhombomere 7). We observed slight decreases in neuronal activity within the forebrain (in regions olfactory bulb, subpallium, pallium) within the diencephalon (habenula, retinal arborization fields) and within the rhombencephalon (inferior olive). These changes in neuronal activity are Cx36-independent and are likely off-target effects. This indicates that, less the off-target effects on neuronal activity we identified in *cx35*.*5-/-* animals (Supplementary Figure 1), the increases in activity we observed in the mefloquine treated fish (hypothalamus, retinal arborization fields, pre-tectum, and subpallium (Figure 5K) are likely due to true increases in activity following only acute blockade of Cx36. For a complete list of regions changed, see Supplementary Table 4.

## DISCUSSION

The goal of this study was to understand the reciprocal relationship between Cx36 and neuronal hyperactivity on a brain-wide scale. We utilized MAP-mapping to quantify neuronal activity and protein expression across the *whole-brain*, which has not been possible using other models. Through this, we characterized the complex nature of this relationship and its dependence on many factors including brain region, drug dose, and exposure time. We found that chronic deficiency of the Cx36 protein in the *cx35*.*5* mutants altered susceptibility to PTZ-induced neuronal hyperactivity in a region-specific manner. We also developed a whole-brain quantification method for Cx36 expression and found that PTZ exposure results in an acute decrease in the expression of Cx36, followed by recovery and overexpression of the protein. Finally, we observed that acute knockdown of the functionality of Cx36 by mefloquine resulted in a broad increase in the susceptibility to PTZ induced hyperactivity. Taken together, these results suggest that Cx36 acts to prevent hyperactivity within the brain, and that loss of Cx36 protein, both acute (perhaps due to previous hyperactivity) and chronic, results in an increase in susceptibility to hyperactivity. As such, preservation of Cx36 expression may serve as a viable therapeutic target in the treatment of diseases such as epilepsy.

### PTZ exerts brain-wide and region-specific effects

We generated dose-varying whole-brain activity maps for PTZ in *cx35*.*5-/-* and wild-type fish. Using the zebrafish model we discovered regions affected by PTZ that were not examined in previous studies. This is important because previous studies in mammalian systems were restricted to the hippocampus. Only 60% of epilepsy cases are characterized by hippocampal sclerosis, with 0.003% of those patients suffering from drug-resistant epilepsy (Asadi-Pooya, Stewart, Abrams, & Sharan, 2017). In all forms of epilepsy, however, approximately 30% of cases are drug-resistant (Kwan & Brodie, 2000). It is therefore imperative to look beyond the hippocampus to address this unmet need.

We did see a slight increase in activity in the pallium at all concentrations of PTZ (analogous to the hippocampus) (Cheng, Jesuthasan, & Penney, 2014) (Figure 1), but it was not the largest increase we observed. We showed a dose-varying dependent increase in activity after administration of PTZ (Figure 1) with larger increases in regions associated with hormone release, and production, as well as executive functioning. These results stress the lack of generalizability of results across brain regions, and the need for expanded inquiry when examining neuronal hyperactivity.

While we were able to examine a greater number of brain regions than previous studies, we sacrificed temporal resolution (achieved with Ca2+ imaging and EEG). However, these results can be used to inform which brain regions should be investigated using methods that allow for greater temporal resolution. In addition to discovering new brain regions affected by PTZ, we were able to elucidate the dose-varying effects of PTZ in a way that was previously unachievable by examining the whole brain. Previous studies, using live calcium imaging, observed increases in neuronal activity and synchronicity after PTZ administration, with differential recruitment of different brain regions (Diaz Verdugo et al., 2019; Liu & Baraban, 2019). They observed increases in neuronal activity originating in the pallium and traveling to the hindbrain (Liu & Baraban, 2019). Additionally, they observed significant increases in neuronal connectivity in each of the regions observed (Diaz Verdugo et al., 2019). Our results show similar effects of PTZ on brain activity in similar regions, but we were able to identify additional brain regions than was previously possible (Diaz Verdugo et al., 2019; Liu & Baraban, 2019). This demonstrates the importance of identifying brain-wide region-specific effects when examining hyperactivity. Taken together, these results illustrate the unique dose-varying whole-brain effects of PTZ that can be expanded upon in future work.

### Cx36 knockdown causes region-specific changes in hyperactivity following PTZ administration

In addition to characterizing the effect of PTZ on whole-brain activity in wild-type animals, we gained insight into the drug’s effects in *cx35*.*5-/-* zebrafish. We found neuronal activity differences in *cx35*.*5-/-* compared to wild-type following high concentrations of PTZ (Figure 1). We saw increases in regions identified in our PTZ dose-response experiment, indicating more severe increases in neuronal hyperactivity following the administration of PTZ in those regions (Figure 1). These results are consistent with previous behavior work by Jacobson, et. al, 2010, which showed that in Cx36 mutant mice, PTZ administration resulted in more severe seizure-associated behaviors than their wild-type counterparts (Jacobson et al., 2010), but also provides more information relating to the severity of neuronal hyperactivity. In addition to activity increases, we observed significant decreases in neuronal activity at 10mM PTZ concentrations. These decreases were observed in the rhombencephalon, specifically in regions that show high Cx36 expression (Figure 3A) and rely on Cx36 for synchronous firing (inferior olive, Mauthner cells) (Bazzigaluppi et al., 2017; Flores et al., 2012; Yao et al., 2014). These results are important, as it is the first study to show regional differences in neuronal activity between Cx36-deficient and wild-type animals, which indicates the lack of generalizability from region to region within the brain when examining connexin proteins.

### PTZ induced hyperactivity causes a regionally-specific decrease in Cx36 expression

To further understand the relationship between Cx36 and hyperactivity, we asked the reciprocal question: how does hyperactivity affect Cx36? Similar to the seizure susceptibility studies, work to identify this relationship has remained conflicting (Laura et al., 2015; Motaghi et al., 2017; Söhl et al., 2000; X. Wu et al., 2017). Previous approaches used to address this question (e.g., qPCR, western blot) lacked the necessary spatial resolution to determine if the effects of hyperactivity on Cx36 vary based on the brain region. To address these shortcomings, we developed a novel method for quantifying the whole-brain expression of the Cx36 protein, using antibody staining in conjunction with a modified MAP-mapping technique (Figure 3). We were, therefore, able to determine that there are regional and exposure time differences in the reduction of Cx36 in response to seizure induction using PTZ. Specifically, we saw reductions in a region-specific manner after exposure to PTZ for 30 minutes, and those reductions were greater after 1 hour of PTZ exposure (Figure 4). Therefore, we have determined that results found in one region of the brain and that PTZ exerts region-specific effects on Cx36.

### Reduction in Cx36 expression following hyperactivity is acute and recovers over time

After observing a decrease in Cx36 expression following exposure to PTZ, we measured the temporal patterns of this change. We found that the change in Cx36 expression was acute: it occurred within the first hour of PTZ exposure and was almost fully recovered by 3 hours (Figure 4C-D). The recovery was then overshot, and the protein was overexpressed in the optic tectum and cerebellum as well as other brain regions, and this overexpression was maintained 24 hours later (Figure 4E-F). Because the reduction was not caused by an increase in cell death (Figure 4G), this effect is likely due to an increase in endocytosis and degradation of the Cx36 protein. Various studies have shown that activity-dependent modulation of Cx36 proteins exists (Haas, Greenwald, & Pereda, 2016; Smith & Pereda, 2003) and endocytosis is a likely mechanism by which this can occur (Flores et al., 2012).

### Acute reduction in Cx36 functionality leaves organisms more susceptible to PTZ induced hyperactivity

To solidify the relationship between hyperactivity and Cx36, we studied how acute blockade of Cx36 affects susceptibility to hyperactivity. Is the reduction in Cx36 after PTZ exposure adaptive, maladaptive, or inconsequential? To answer this question, we utilized the Cx36 specific blocking drug mefloquine and expose mefloquine treated and untreated fish to PTZ to observe differences. Mefloquine is an anti-malarial drug that selectively blocks Cx36 and Cx50. Previous studies utilized quinine which has more off-target effects. It is hypothesized that mefloquine blocks Cx36 by binding to the inside of the pore, preventing the flow of ions through that pore (Harris & Locke, 2008). We found a significant increase in neuronal hyperactivity following treatment with PTZ in the mefloquine treated fish compared to control (Figure 4). This result indicates a reduction in Cx36 in all cases (acute and chronic) is detrimental and leads to an altered severity of hyperactivity.

At moderate doses (6-25 μM), mefloquine can exhibit off-target effects of varying degrees (Caridha et al., 2008; Harris & Locke, 2008; McArdle, Sellin, Coakley, Potian, & Hognason, 2006). To control for off-target effects of mefloquine, we treated *cx35*.*5-/-* fish with mefloquine and quantified changes in neuronal activity both at rest (in embryo medium) and after PTZ (5 mM). We observed major decreases in neuronal activity within the forebrain and a slight decrease in the rhombencephalon in both conditions. Additionally, we observed a slight increase in neuronal activity in the rhombencephalon which was exacerbated slightly by PTZ (Supplementary Figure 1). We attribute these effects to off-target effects of mefloquine, while the changes in PTZ sensitivity caused by mefloquine in other ideas are more likely to be caused by Cx36 blockade.

The increase in neuronal hyperactivity following treatment with PTZ in the mefloquine treated fish compared to control (Figure 4) is greater than the *cx35*.*5-/-* to wild-type comparison (Figure 1). This may be due to genetic compensation in the *cx35*.*5* mutants resulting from the lack of Cx36 from birth. It is also possible that acute reduction of Cx36 is more detrimental than chronic knock down, meaning the acute reduction in Cx36 expression after hyperactivity, would be more detrimental than chronic knock-down. This difference may then also account for conflicting evidence in the field comparing chronic and acute knockdown of Cx36 (Gajda et al., 2005; Jacobson et al., 2010; Voss et al., 2010b). Taken together, these results suggest that the prevention of the loss of Cx36 function may prove to be a useful target for treating diseases of hyperactivity.

### Cx36 is a contributing factor regulating the brains response to hyperactivity

A plausible clinical application of Cx36-targeted therapeutics is in Juvenile Myoclonic Epilepsy (JME). Individuals with JME have a higher likelihood of harboring a specific intronic SNP in the *Cx36* gene (Hempelmann, Heils, & Sander, 2006; Mas et al., 2004). This SNP has been hypothesized to affect splicing enhancers of the gene, therefore affecting the translation of the protein (Mas et al., 2004). While Cx36 may not be the only cause for diseases like JME, it may be a contributing factor. Based on our results, loss of Cx36 makes an individual more susceptible to other factors leading to hyperactivity, increasing the severity of hyperactivity (Figure 1, 6), therefore, the rescue of Cx36 expression may reduce the severity of hyperactivity. This is particularly relevant as Cx36 expression is highest during development and decreases over time (Belousov & Fontes, 2013) and JME first appears in children and adolescents. This is clinically relevant as approximately 15% of cases of JME are drug-resistant, with no known pharmacotherapy (Martin et al., 2019).

Our work demonstrates that Cx36 is an important factor preventing hyperactivity in the brain and that loss of the protein is detrimental to that process. We were able to determine where in the brain we see effects in addition to when those changes occur. This work provides a basis for better understanding the dynamics of Cx36 and hyperactivity.

## Acknowledgments

This work was supported by funding from the Commonwealth Research Commercialization Fund (ER14S-001-LS to Y.A.P.) and Virginia Tech. We thank the animal care staff at Virginia Tech for animal husbandry and Dr. Adam Miller for the *cx35*.*5* mutant zebrafish. We are also appreciative of Dr. Susan Campbell and Dr. James Smyth for their helpful suggestions.

## Figure Legends

**Supplementary Figure 1.**
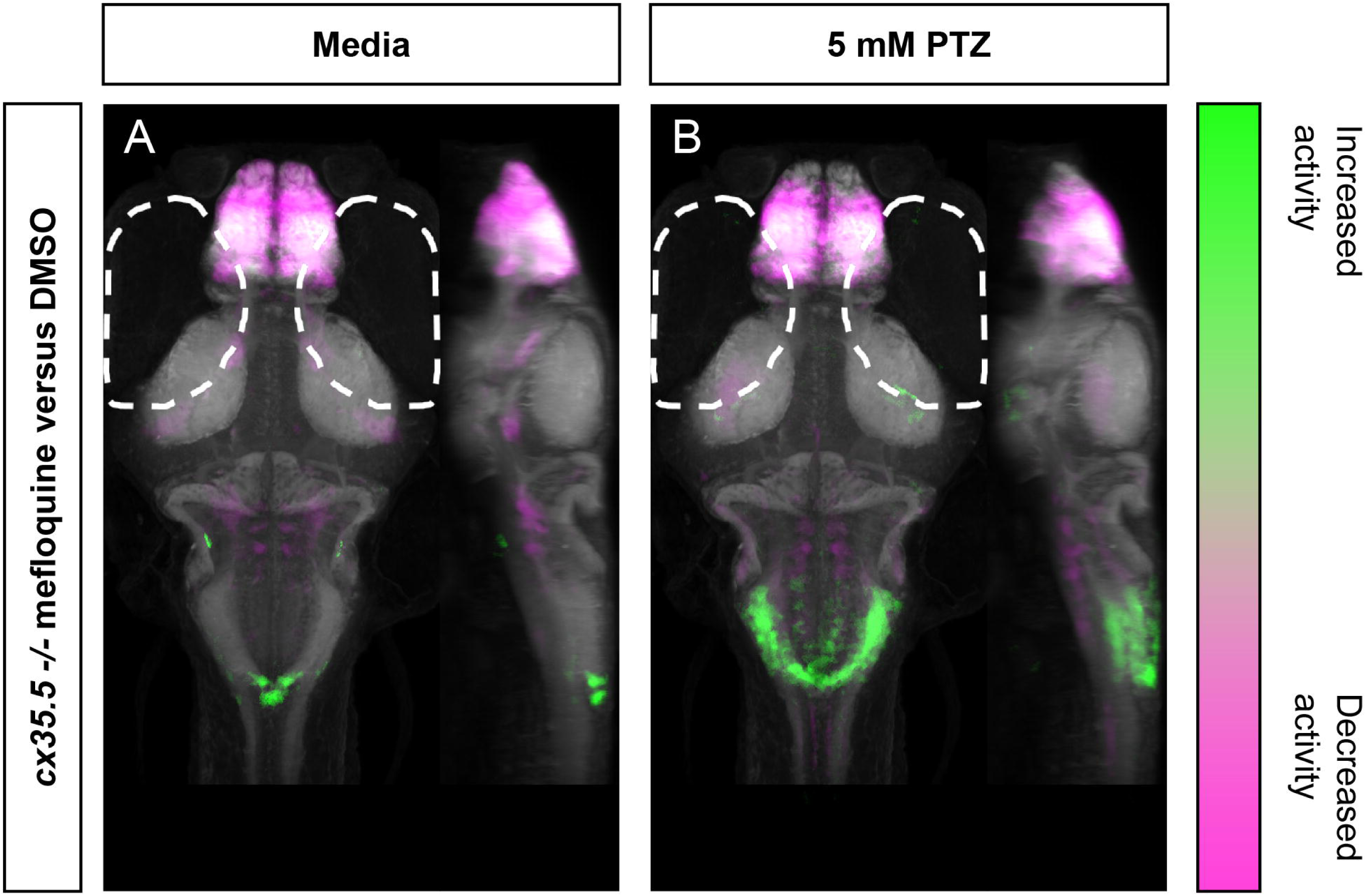
Whole-brain activity map showing off-target effects of mefloquine using *cx35*.*5-/-* treated with mefloquine. Dorsal and lateral view of zebrafish larvae brain. Images show pixels with significantly increased pERK/tERK ratio in A) Embryo medium and mefloquine treated compared to DMSO treated larvae. (*cx35*.*5-/-* mefloquine n=9, *cx35*.*5-/-* DMSO n=11) and B) 5 mM PTZ and mefloquine treated compared to DMSO treated larvae (*cx35*.*5-/-* mefloquine n=11, *cx35*.*5-/-* DMSO n=11). For all regions p < 0.005

